# Smoke exposure triggers rapid transcriptional changes and reveals glycosyltransferases linked to smoke taint in ripening grape berries

**DOI:** 10.1101/2025.02.11.637658

**Authors:** Manon Paineau, Chen Liang, Noe Cochetel, Shivani, Rosa Figueroa-Balderas, Roger Thilmony, Arran Rumbaugh, Dario Cantu

## Abstract

With the increasing frequency of wildfires, vineyards are more often exposed to smoke, resulting in a higher risk of smoke taint in wine. This taint imparts undesirable “smoky” and “ashy” aromas and causes significant economic losses. Smoke-derived volatile phenols (VPs) in grape berries, metabolized into stable, non-volatile glycoconjugates via glycosyltransferases (GTs), underlie smoke taint formation. Here, we present the dwarf grapevine cultivar Pixie as a model system for smoke taint research. We generated a phased, telomere-to-telomere diploid genome using HiFi long-read sequencing, along with an expression atlas spanning 30 replicated samples across various tissues and developmental stages. We also constructed a second telomere-to-telomere assembly for Pinot Noir to assess *GT* expression during berry ripening in the field. Controlled smoke-exposure experiments profiled *GT* expression and enabled transcriptome-wide analyses at multiple time points. We measured VPs in both smoke-exposed and control samples, revealing 12 *GT1* genes significantly upregulated by smoke, named *Vitis vinifera smoke-inducible UGTs* (*VviSIUGTs*). Their expression peaked one day post-exposure and declined by day three, mirroring glycoside accumulation dynamics. Gene co-expression network analyses highlighted stress-responsive modules enriched in these *VviSIUGTs*, while transcription factor binding site analyses pinpointed stress-related regulatory elements in their promoters. These findings provide new insights into the molecular basis of smoke taint and identify potential targets for breeding and biotechnological interventions aimed at developing grapevine cultivars with reduced susceptibility to smoke damage.

## Introduction

Wildfires frequently occur in regions such as North America, Australia, South Africa, and the Mediterranean. Rising temperatures, strengthening winds, and prolonging droughts have intensified these events (Abbass *et al*., 2022). Many vineyards are located in Mediterranean-type climates, where both prescribed burns and unplanned fires are common. Although flames can directly damage vines, smoke exposure poses a significant economic risk, often rendering grapes and wine unmarketable. To put this into perspective, in 2020, severe wildfires burned over 10 million acres in the United States equating to an estimated $3.7 billion in economic losses for the grape and wine industry. Grapes rejected by wineries hit a record high in 2020, where $601 million of California grapes were not harvested due to potential smoke impact (Kropp and De Andrade, 2022).

When smoke drifts into vineyards, smoke-derived volatile phenols (VPs)—byproducts of plant lignin thermal degradation, such as guaiacol, 4-methylguaiacol, syringols, and cresols—are absorbed by the grapes (Summerson *et al*., 2021). Once absorbed, these VPs are metabolized into non-volatile glycoconjugates, that, upon fermentation, lead to smoke taint in wine (Hayasaka *et al*., 2010). The conversion of the VPs into their stable glycosidic form suggests the involvement of glycosyltransferases (GTs). These enzymes are currently classified into 137 families in the CAZyme database based on sequence similarity, catalytic activity, and conserved sequence motifs (Campbell *et al*., 1997; Ross *et al*., 2001). Among them, family 1 glycosyltransferases (GT1s) is the largest group and, in plants, it predominantly includes uridine diphosphate-dependent glycosyltransferases (UGTs) (Louveau and Osbourn, 2019). UGTs transfer sugars from nucleotide-activated donors to acceptor molecules (Hughes and Hughes, 1994; Lairson *et al*., 2008; Osmani *et al*., 2008; Rahimi *et al*., 2019). Given their broad role in plant stress responses and environmental adaptation (Gharabli *et al*., 2023), UGTs are strong candidates for mediating glycosylation of smoke-derived VPs. Understanding the activity of GT1 enzymes, particularly UGTs, is therefore critical for elucidating the metabolic pathways for VP glycosylation in smoke-exposed grapevines.

The chemical basis of smoke taint in wine has been extensively studied (Summerson *et al*., 2021), yet relatively few studies have examined the genetic mechanisms involved, particularly the role of grape GTs. Prior findings suggest GT involvement: for example, *UGT72B27*, a resveratrol glycosyltransferase that can conjugate smoke-derived phenolics, is highly expressed in grapevine leaves and berries (Härtl *et al*., 2017). Similarly, five *GT* genes were identified as upregulated in smoke-exposed Chardonnay and Shiraz, indicating that both cultivar and tissue type influence *GT* gene expression patterns (van der Hulst, 2018). More recently, smoke exposure was found to upregulate genes associated with detoxification and cell wall reinforcement, with GTs playing a central role in these responses (Hewitt *et al*., 2024). In this study, we profiled *GT1* gene expression in grape berries following smoke exposure under controlled conditions, thereby eliminating field-related confounding factors. We employed the Pixie dwarf grapevine mutant, derived from the L1 cell layer of Pinot Meunier (Boss and Thomas, 2002; Cousins, 2012). Pixie vines remain small, produce fruit continuously, and thrive under greenhouse conditions, making them an ideal system for reproducible experiments on the effects of smoke exposure in grapes. We assembled a telomere-to-telomere diploid Pixie genome and annotated its *GT1* genes by referencing a comprehensive library of 4,205 *GT1* genes identified across eight diploid grapevine genomes. Subsequently, we developed a detailed gene expression atlas covering three replicates of ten samples from multiple tissues and developmental stages, profiled *GT* expression over the course of berry ripening, and conducted a targeted smoke-exposure time-course experiment to examine *GT* expression dynamics in response to smoke. By integrating transcriptome profiles, gene co-expression networks, and VP accumulation data, we uncovered a set of twelve smoke-inducible *GT1* genes. These candidate genes form part of specific co-regulated networks enriched in stress-responsive transcription factors. Together, these findings provide a more comprehensive understanding of the genetic networks underlying berry response to smoke and highlight actionable targets for breeding and biotechnological strategies.

## Materials and methods

### DNA extraction and HiFi sequencing libraries for genome assemblies

Pixie clone 01 and Pinot Noir clone 123 are maintained by the Foundation Plant Services at the University of California, Davis. High-molecular-weight genomic DNA (gDNA) was isolated from young leaves using the method described in (Chin *et al*., 2016). DNA purity was evaluated with a Nanodrop 2000 spectrophotometer (Thermo Scientific, IL, USA), DNA quantity with the DNA High Sensitivity kit on a Qubit 2.0 Fluorometer (Life Technologies, CA, USA), and DNA integrity by Agilent FEMTO Pulse (Agilent, CA, USA). SMRTbell library from PN123 was prepared and sequenced as described in (Cochetel *et al*., 2023). HiFi sequencing library from Pixie FPS01 was prepared using the SMRTbell Express Template Prep Kit 2.0 (Pacific Biosciences, CA, USA) following the manufacturer’s instructions. Library was size selected using 3.3X v/v of 35 % Ampure PB beads (Pacific Biosciences, CA, USA). The size-selected SMRTbell library was sequenced on a PacBio Sequel II platform using a V2.0 chemistry (DNA Technology Core Facility, University of California, Davis, CA, USA).

### RNA extraction, library preparation, and sequencing

The Pixie plants used in the experiment have been kept in a greenhouse since 2019. Prior to the experiment they were maintained to only have 2 clusters per shoot. Plants were watered 3 times a day for 10-minutes increments. Growing temperatures were as follow: day 75 to 85F (24 to 29C), night 60 to 71F (16 to 22C). A total of 30 samples were collected for the Pixie expression atlas. Berry samples included three developmental stages: green berry (hard, green, pre-veraison), harvest berry (colored, soft, ripe), and post-harvest berry (very soft, overripe). Leaf samples were from three stages: young leaf (first two leaves near the shoot tip), green leaf (fully developed, healthy leaves), and senescent leaf (older leaves with brown patches). Additional tissues included root (clean samples from in vitro growth), stem (young, green stems), buds (secondary dormant buds) and flower (entire flowers post-cap fall, excluding cap and pedicel). RNA was extracted from these samples using the method described in (Rapicavoli *et al*., 2018) with the exception of the deseeded berries that was isolated using a Cetyltrimethyl Ammonium Bromide (CTAB)-based extraction protocol as described in (Blanco-Ulate *et al*., 2013). RNA purity and quantity were evaluated using a Nanodrop 2000 spectrophotometer (Thermo Scientific, Hanover Park, IL, USA) and a Qubit 2.0 Fluorometer (Life Technologies, Carlsbad, CA, USA). RNAseq libraries were prepared using the Illumina TruSeq RNA Sample Preparation Kit v2 (Illumina, CA, USA) following Illumina’s low-throughput protocol. The library was evaluated for quantity and quality using the High Sensitivity chip on an Agilent 2100 Bioanalyzer (Agilent Technologies, CA, USA) and sequenced on the Elements Bio Aviti platform at PE80 (DNA Technology Core Facility, University of California, Davis, CA, USA).

### Pixie and Pinot Noir genome assembly, gene prediction and functional annotation

The HiFi reads of Pixie were assembled using a two-step procedure involving Hifiasm v.0.16.1- r37412 (Cheng *et al*., 2021, 2022) and the HaploSync tool suite v1.0 (Minio *et al*., 2022). To obtain the least fragmented draft genome assembly, various combinations of parameters were tested with HiFiasm to determine the optimal settings for constructing a high-quality assembly. These parameters were evaluated based on the expected haplotype size, contig length, and fragmentation of the draft assembly. A total of 326 diploid assemblies were generated to identify the best configuration. The selected parameters (a=2, k=61, w=61, f=25, r=4, s=0.6, D=3, N=25, n=13, m=10,000,000) produced a draft assembly of 497.4 Mbp with 208 contigs for haplotype 1. This assembly had an average contig length of 2.4 Mbp and an N50 of 20.5 Mbp (L50 = 10). Haplotype 2 reached 506.2 Mbp with 85 contigs, an average contig length of 5.9 Mbp, and an N50 of 21.0 Mbp (L50 = 10). We then constructed scaffolded, phased, chromosome-scale pseudomolecules for both haplotypes (Hap1 and Hap2; **Table 1**) using the consensus grape genetic map (Zou et al., 2020) and HaploSync v1.0 (Minio *et al*., 2022). Two iterations of HaploSplit and two iterations of HaploFill were required to minimize fragmentation and achieve the final assembly.

**Table 1.** Assembly statistics of the Pixie’s genome.

| HiFiasm assembly results | Haplotype 1 | Haplotype 2 |  |
| --- | --- | --- | --- |
| Total contig length (bp) | 497,387,120 | 506,190,299 |  |
| Contig N50 Length (bp) | 20,464,270 | 20,999,202 |  |
| Contig N50 index | 10 | 10 |  |
| Maximum contig length (bp) | 36,866,322 | 37,596,281 |  |
| Haplosync pseudomolecule reconstruction results | Haplotype 1 | Haplotype 2 | Unplaced |
| Total pseudomolecule length (bp)* | 490,897,489 | 496,247,121 | 17,823,266 |
| N. of pseudomolecules* | 19 | 19 | 224 |
| N. of pseudomolecules with 2 terminal telomeric repeats | 15 | 17 | - |
| N. of pseudomolecules with only 1 terminal telomeric repeat | 4 | 2 | - |
| GC content (%) | 35 | 35 | 45 |
| Repeats content (%) | 58 | 58 | 73 |
| N. of protein coding-genes | 24,524 | 24,605 | 476 |
| Complete BUSCO (%) | 99.1 | 98.8 | 1.4 |

\* Unplaced = N. and total size of unplaced contigs

Pinot Noir (clone FPS 123) CLR PacBio reads were assembled using a two-step procedure involving the FALCON-Unzip pipeline (v.2017.06.28-18.01; Chin et al., 2016) and the HaploSync tool suite v1.0 (Minio *et al*., 2022). The scaffolding process was performed as described in (Minio *et al*., 2019). The customized FALCON-Unzip pipeline is available at https://github.com/andreaminio/FalconUnzip-DClab. The draft assembly (**Table S1**) consisted of 1,490 primary contigs with a combined size of 618.1 Mbp and 2,532 haplotigs with a total size of 245.2 Mbp. The primary contigs and their associated haplotigs were then phased and assembled at the chromosome level using the consensus grape genetic map (Zou et al., 2020) and HaploSync v1.0 (Minio *et al*., 2022). A total of eight iterations of HaploSplit and five iterations of HaploFill were required to minimize fragmentation and achieve the final assembly. The final assembly of Pinot Noir reached a total length of 442.5 Mbp for haplotype 1 and 422.6 Mbp for haplotype 2. Less than 10% of the diploid assembly (80.0 Mbp; 1,245 contigs) remained unplaced. For both genomes, gene structural annotations were predicted using the procedures described in https://github.com/andreaminio/AnnotationPipeline-EVM_based-DClab21. The telomere repeat units were analyzed using TIDK (v.0.2.63, Brown *et al*., 2025). The genomes were explored with *tidk explore*, focusing on 1% of the chromosome ends, with telomeric repeat units ranging from 5 to 12 base pairs in length and a threshold of 2 sequential repetitions. The telomeric repeat sequence TTTAGGG was then searched using *tidk search*. The results were visualized with *tidk* plot (**Figure S1**).

### Comparison of genome assemblies

Synteny analysis was performed using MCScanX (version 1.0.0, Wang *et al*., 2012), with a minimum of 10 genes required to define a collinear block and up to 10 gene gaps allowed within blocks. Whole proteomes of Pixie, Pinot Noir, and PN40023 were compared using blastp (version 2.12.0+, Camacho *et al*., 2009)) (parameters: -e 1e-10, -b 5, -v 5). Synteny visualization was carried out with SynVisio (Bandi and Gutwin, 2020). Pairwise chromosome alignments were generated using NUCmer from MUMmer v.4.0.0 (Marçais *et al*., 2018), with a minimum match length of 3000 (option -l) filtered using nucmer --filter. Visualizations were created in R using the alignment data from mummer show-coords, with -L 5000 set for improved figure clarity. Single nucleotide polymorphisms (SNPs) and small insertions and deletions (InDels) were called with show-snps, while structural variants (SVs) were detected with show-diff.

### Glycosyltransferase family 1 (*GT1*) identification and annotation

To identify and annotate *GT1* genes in the Pixie genome, we first constructed a GT1 database specific to *V. vinifera*. This was achieved using Hidden Markov Models (HMMs) based on GT1 protein domain composition, leveraging the CAZy (http://www.cazy.org/) and Pfam (http://pfam.xfam.org/) databases. Eight diploid grapevine genomes (**Table S1**) were screened using *hmmscan* (from HMMER version 3.3.2, http://hmmer.org/), with a threshold set to exclude domains with an e-value greater than 1e-7. This screening identified 11,238 GTs from Pfam and 24,264 from CAZy. Focusing on GTs containing the PF00201 domain or annotated as GT1, we identified 9,394 *GT* genes, including 4,374 with the PF00201 domain and 5,050 annotated as *GT1*. Subsequently, we developed grapevine-specific HMM models by clustering *GT1* sequences based on their size and similarity using CD-HIT2. One representative model was generated for each cluster to ensure optimal precision, resulting in 28 models for PF00201 and 44 models for CAZy. This refined approach identified a total of 4,205 *GT* genes across the eight genomes. To annotate *GT1*s in the Pixie genome, we mapped our *V. vinifera GT1* collection onto the newly assembled Pixie genome using GMAP (version 2019-09-12, Wu and Watanabe, 2005) and filtered the results keeping a coverage and similarity threshold of 0.95. A total of 514 GT1s were identified in Pixie, with 263 found in the first haplotype and 241 in the second.

### Phylogenetic analysis

The multiple sequence alignment (MSA) was performed using MAFFT (v7.511, Katoh *et al*., 2002). To retain only the conserved regions of the proteins, the MSA was trimmed using Gblocks (v0.91b, Talavera and Castresana, 2007). Parameters included a minimum number of sequences for a conserved or flank position corresponding to half the number of aligned proteins, a maximum of 25 contiguous non-conserved positions, a minimum block length of 3, and allowed gaps in positions present in at least half of the sequences. Four proteins (*VITVvi_vPixieFPS01_v1.0.Hap2.chr16.ver1.0.g446810.t01*, *VITVvi_vPixieFPS01_v1.0.Hap1.chr16.ver1.0.g194610.t01*, *VITVvi_vPixieFPS01_v1.0.Hap2.chr16.ver1.0.g439860.t01*, and *VITVvi_vPixieFPS01_v1.0.Hap2.chr08.ver1.0.g350410.t01*) were removed from the analysis because their PSPG boxes were not conserved, which disrupted the alignment of this critical region. The filtered alignment retained 4% of the sequences, distributed across 17 blocks, for the alignment of the 480 Pixie UGTs and 2% of the sequences, distributed across 16 blocks, for the alignment of the 480 Pixie UGTs with 133 *Arabidopsis thaliana* UGTs retrieved on https://pugtdb.biodesign.ac.cn/family. Maximum-likelihood phylogenetic trees were inferred using RAxML-NG (version 0.9.0, Kozlov *et al*., 2019) with the best-fit models identified by ModelTest-NG (Darriba *et al*., 2020): LG4M+G4 for Pixie UGTs and JTT+G4 for Pixie and *A. thaliana* UGTs. Each tree was built with 700 bootstrap replicates and five starting parsimony trees. The trees were visualized using the R package ggtree (version 3.6.2, Yu *et al*., 2017). Analysis on R were executed with R software version 4.2.2 (R Core Team 2022).

### Protein structure analysis

Structural prediction of the candidate UGTs’ structure was performed using ALPHAFOLD3 (version 3.0.1, Abramson *et al*., 2024) using default settings. Visualization and superimposition of predicted structures were performed on UCSF Chimera X 1.5 (Goddard *et al*., 2018). The predicted structure of UGT72B1 and UGT74F2 were retrieved from the EBI ALPHAFOLD2 database (https://alphafold.ebi.ac.uk/entry/Q9M156) and (https://alphafold.ebi.ac.uk/entry/O22822), respectively.

### Smoke exposure experiment

Entire grape clusters from the Pixie vine were exposed to smoke in two bench-top desiccation chambers: one containing smoke and the other serving as a control without smoke. Approximately seven days post-veraison, the grape clusters were sterilized prior to the experiment using a 200- ppm bleach solution. The experiment was performed twice. In Experiment A, all plants were grown in the Davis greenhouse. A total of 14 clusters were collected, with 2 to 3 clusters per plant. These clusters were evenly divided between the smoke and control treatment groups to minimize variation among individual plants. In Experiment B, eight Pixie vine clusters were used, with the plants grown in two different greenhouse locations, Davis and Albany, both in California, USA. To account for potential location-based effects, clusters from the two greenhouses were evenly split between the smoke and control treatment groups. The two bench-top desiccation chambers were modified to accommodate smoke from a wood burner, saturating the chamber with smoke (**Figure S2**). A small aperture allowed smoke to enter the chamber, while a second opening let air escape, drawing in more smoke. The smoke was generated by burning oak wood chips, with 2 grams consumed over the two-hour exposure period, delivered at a rate of 0.25 grams every 15 minutes. After two hours of exposure, both openings were sealed, allowing for an additional one-hour incubation of the clusters in the residual smoke. During the three-hour experiment, air quality metrics—temperature, humidity, and volatile compounds—were continuously monitored using an air quality monitor (Wave Mini, Airthings, Oslo, Norway). After incubation, the clusters were placed at room temperature in a homemade ’resting chamber’ constructed from plastic sheeting around a PVC frame (**Figure S2E**). A tray of water was added for humidity control, and a small fan directed airflow from the control grapes to the smoke-exposed grapes to prevent contamination of the control grapes. Berry samples for RNA extraction were collected at two time points: 24 hours (T1) and 72 hours post-incubation (T3), resulting in 28 samples in Experiment A and 16 samples in Experiment B. For each sample, 10 to 12 berries were taken per cluster, ensuring uniform sampling without damaging any remaining berries. The seeds were carefully removed, and only the skin and pulp were retained. The samples were immediately flash-frozen in liquid nitrogen and stored at -80°C until RNA-seq and volatile phenol extractions were performed.

### Volatile phenolics (VP) analysis

Standards of VPs were purchased from Sigma-Aldrich, and isotopically labeled internal standards (ISTD) were obtained from CDN Isotopes. HPLC-grade solvents (methanol, hexane and ethyl acetate) were purchased from Sigma-Aldrich. The VP extraction protocol followed the procedures described by (Noestheden *et al*., 2017) with slight modifications. One gram of thawed homogenized Pixie berries was weighed into a 15 mL centrifuge tube, and 4 g of ultrapure water was added. The diluted homogenized grape (1:5) was spiked with ISTD (20 ng/g), and 2 mL hexane/ethyl acetate (1:1, v/v) was added for extraction. Spike-recovery samples were prepared to validate the calibration. Analyses were performed on an Agilent 8890 GC coupled with a 7000D Triple Quad. Instrument parameters and data acquisition followed the method described by (Rumbaugh *et al*., 2025).

### Glycosylated phenol (Gly-VP) analysis

Gly-VP standards (guaiacol gentiobioside, *p*-cresol rutinoside, 4-methylsyringol gentiobioside) and isotopically labeled internal standards (guaiacol gentiobioside-d_3_, *p*-cresol rutinoside-d_7_, 4- methylsyringol gentiobioside-d_6_) were purchased from LGC standards. LC-MS grade solvents (acetonitrile, methanol) were obtained from Sigma Aldrich. Sample preparation followed the procedure reported by (Caffrey *et al*., 2019). Tandem mass spectrometry was performed on an Agilent 6460 triple quadrupole spectrometer with an Agilent Jetstream electrospray source. Instrument settings and data acquisition followed the method by (Lim *et al*., 2024). Guaiacol glucoside, guaiacol rutinoside and 4-methylguaiacol rutinoside were quantified as guaiacol gentiobioside equivalent. Phenol rutinoside was quantified as cresol rutinoside equivalent. Syringol gentiobioside was quantified as 4-methylsyringol gentiobioside equivalent.

### RNAseq data analysis

RNAseq data were processed using the Nextflow pipeline nf-core/rnaseq (v3.12.0) (https://github.com/nf-core/rnaseq/tree/3.12.0; 10.5281/zenodo.1400710). Briefly, adaptor and quality trimming were performed with *TrimGalore* (version 0.6.7, (Krueger, 2025)), and reads were aligned to the Pixie transcriptome using *Salmon* (version 1.10.1, (Patro *et al*., 2017). Quantification was conducted with Salmon using the parameters --seqBias and --posBias. The quantification results were imported, and TPM values were extracted using the R package *tximport* (version 1.24.0, (Soneson *et al*., 2015)). Differential gene expression analysis was performed with the R package *DESeq2* v.1.38.2 (Love *et al*., 2014). We considered a gene as differentially expressed if its log2(FoldChange) is greater than 1 (for upregulated genes) or less than -1 (for downregulated genes), with an adjusted p-value (Padj) lower than 0.05. The heatmap of the Transcripts Per Million (TPM) values was visualized using the *pheatmap* package (version 1.0.12, Kolde, 2019) and the Volcano plot to visualize the log2 fold change was visualized using the R package *EnhancedVolcano* (version 1.16.0, Blighe, 2018). Analyses in R were executed with R software version 4.2.2 (R Core Team 2022).

For the expression atlas, the visualization of transcript expression, transformed to log10(TPM+1) for comparison purposes, was performed using the R package *pheatmap* (version 1.0.12, Kolde, 2019). A user-friendly Expression Atlas App, available on https://grapegenomics.com/pages/VvPixie/Pixie_Atlas_Site/atlas.php, was developed using the R package *Shiny* (version 1.9.0), enabling users to explore the expression of a list of transcripts across various Pixie tissues.

### Gene ontology enrichment analysis

Gene ontology enrichment was performed using the R package TopGo (version 2.50.0, Alexa and Rahnenfuhrer, 2024). Enrichment analysis was conducted using Fisher’s exact tests, with a GO term considered significantly enriched if the adjusted p-value (Bonferroni method) was less than 0.05.

### Weighted gene co-expression network analysis

We conducted a weighted gene co-expression network analysis (WGCNA) using the software *WGCNA* package (version 1.73) in R (Langfelder and Horvath, 2008) on 41,732 transcripts that met the threshold of at least one sample with TPM ≥ 1, from the original set of ∼70,000 transcripts. A Topological Overlap Matrix (TOM) was generated using the ‘TOMsimilarityFromExpr’ function with a signed hybrid network type and a soft-thresholding power of 6, determined via the ‘pickSoftThreshold’ function. Modules were detected by hierarchical clustering of 1-TOM (dissTOM) with average linkage, and the dynamic tree cut method (‘cutreeDynamic’ function) with a minimum module size of 50. To analyze module-trait associations, we calculated Pearson correlation coefficients between module eigen-genes (MEs) and experimental traits (experimental variables, free and bounds volatile phenols), deriving p-values with a Student’s t-test adjusted for sample size. The resulting correlations were visualized using the R package *Pheatmap* (version 1.0.12, Kolde, 2019). Module Membership (MM), representing the correlation between a gene and its module eigen-gene, and Gene Significance (GS), measuring gene-trait correlations (trait ’smoke’ has been used), were calculated to identify hub genes. Highly correlated MM and GS values suggested that central genes in each module are key to trait association; hub genes per module were then defined as those with MM > 0.75 and GS > 0.5.

### Transcriptor Factor Binding Site (TFBS) analysis

The promoter sequences of the previously identified hub genes were extracted using a custom Python script to retrieve the 3-kb regions upstream of the transcription start sites (TSS) of each hub gene. Transcription factor binding sites (TFBS) were identified using the R package TFBSTools (version 1.36.0, Tan and Lenhard, 2016) and the JASPAR24 database (Rauluseviciute *et al*., 2024). Enrichment analysis was conducted using Fisher’s exact tests, with TFBS considered significantly enriched if the adjusted p-value (Bonferroni method) was less than 0.05.

### qRT-PCR analysis

Quantitative Reverse Transcriptase PCR (qRT-PCR) was performed using the iTaq Universal SYBR green one step kit (Bio-Rad, CA, USA) for expression analysis of eight *VviSIUTG* genes (primers shown in **Table S5**). The PCR reactions contained 5 µl of 2× SYBR Green Master Mix, 0.125 µl of iScript Reverse Transcriptase (Bio-Rad, Hercules, CA, USA), 0.3 µL each of each forward and reverse primers (10 µM), 4 µL of RNA template (50 ng/µl), and nuclease free water to make up final volume 10 µl. PCR amplification was performed using an Applied Biosystems QuantStudio 3.0 Real-Time PCR thermocycler (Thermo Fisher, CA, USA). The thermal cycling conditions comprised 50°C for 10 minutes and 95°C for 1 min (Stage 1), followed by 40 cycles of denaturation at 95°C for 15 sec and annealing/extension at 60°C for 1 min (Stage 2). Melt curve analysis was performed with a final denaturation step at 95°C for 15 sec, followed by annealing at 60°C for 1 min to confirm the specificity of the amplification. PCR amplification was conducted on three biological and two technical replicates. Gene expression of smoke-treated samples was normalized with control samples, normalized with *VviActin* and *VviGAPDH* reference genes (Reid *et al*., 2006) using the Pfaffl method (Pfaffl, 2001).

### Data availability

The genome assemblies and annotations of Pixie and Pinot Noir can be accessed on Zenodo (Pixie, doi:10.5281/zenodo.14853250; Pinot Noir, doi: 10.5281/zenodo.14853293). Dedicated genome browsers are available on GrapeGenomics.com at https://grapegenomics.com/pages/VvPixie/ for Pixie and https://grapegenomics.com/pages/VvPinNoir/VvPinNoir123/ for Pinot Noir. The Pixie expression atlas is accessible at https://grapegenomics.com/pages/VvPixie/Pixie_Atlas_Site/atlas.php.

## Results

### A telomere-to-telomere diploid genome reference for the Pixie grapevine

To profile the gene expression of Pixie grapevine berries in response to smoke exposure, we began by generating a telomere-to-telomere diploid genome assembly for the Pixie grapevine. Using PacBio’s HiFi long-read sequencing, we produced a total of 25.4 billion bases (bp), distributed across 2,317,432 reads with a median length of 9,994 bp. Using HiFiasm (Cheng *et al*., 2021), we generated and evaluated 326 diploid assemblies to determine the optimal combination of assembly parameters. The best-performing assembly produced two haplotypes: haplotype 1 (hap1) spans 497.4 Mbp across 208 contigs (N50 = 20.5 Mbp), while haplotype 2 (hap2) spans 506.2 Mbp across 85 contigs (N50 = 21.0 Mbp). Both haplotypes were then scaffolded into chromosome-scale pseudomolecules using HaploSync v1.0 and a *Vitis* consensus genetic map (Zou *et al*., 2020; Minio *et al*., 2022). The resulting diploid genome assembly measured 490.9 Mbp for hap1 and 496.2 Mbp for hap2 (**Table 1**). All but six of the pseudomolecules contained terminal telomeric repeat units TTTAGGG at both ends, indicating near telomere-to-telomere (T2T) completeness. The six exceptions (chromosome 5 of hap1, chromosome 15 of both hap1 and hap2, chromosome 17 of both hap1 and hap2 and chromosome 18 hap1) each contained a single terminal telomeric repeat. Quality assessment with the Benchmarking Universal Single-Copy Orthologs (BUSCO, (Simão *et al*., 2015)) tool indicated high completeness, with scores of 99.1% for hap1 and 98.8% for hap2. The unplaced sequences accounted for only 1.7% of the diploid genome assembly and 1.4% of the BUSCO proteins, confirming the overall completeness of the hap1 and hap2 assemblies. These unplaced sequences were predominantly composed of repetitive elements (73%), compared to 58% in the main haplotypes.

The Pixie genome shows strong synteny with both the PN40024_T2T genome and the Pinot Noir FPS123 genome generated as part of this study (**Table S2**), with 83.69% of the genes being collinear across the three assemblies (76.52% with PN40024 and 86.58% with Pinot Noir; **Figure S3**). The genome assemblies exhibit high sequence identity, with over 99% identity observed in both the comparison between Pixie and PN40024_T2T, and the comparison between Pixie and Pinot Noir. The alignments for each chromosome in both comparisons are presented in **Figures S4** and **S5**. A total of 5,463,299 SNPs (143.8 ± 55,513.6 SNPs/chromosome), 998,498 InDels (26,276.3 ± 10,146.6 InDels/chromosome), and 54,394 SVs (1,431.4 ± 559.9 SVs/chromosome) were identified between Pixie and PN40024, while 2,954,598 SNPs (77,752.6 ± 19,959.1 SNPs/chromosome), 736,617 InDels (19,384.7 ± 4,425.1 InDels/chromosome), and 32,041 SVs (843.2 ± 243.9 SVs/chromosome) were identified between Pixie and Pinot Noir, with detailed data for each chromosome provided in **Figure S6**. These results confirm the high degree of genomic conservation and structural integrity among these closely related *V. vinifera* accessions.

### Establishing a specialized *Vitis vinifera* GT1 database for enhanced glycosyltransferase annotation in the Pixie genome

To identify the complete set of *GT1s* in the Pixie reference genome, we first screened for *GTs* in eight annotated diploid *V. vinifera* genomes (**Table S2**), evaluating both haplotypes of each genome. Our initial search identified 11,238 *GTs* showing homology to Pfam (1,386.6 ± 328.5 genes/per cultivar) and 24,255 *GTs* with homology to the CAZy database (2,960.8 ± 757.2 genes/cultivar). Among these, we focused on *GTs* containing the PF00201 Pfam domain, corresponding to UDP-glycosyltransferases (*UGTs*), and those annotated as *GT1s* in the CAZy database. Collectively, we identified 5,889 *GT* genes, comprising 4,374 containing the PF00201 domain (510.3 ± 77.6 genes/cultivar) and 5,019 annotated as *GT1*s (649.4 ± 88.9 genes/cultivar). Of these, 337 *GTs* were uniquely associated with the PF00201 domain, 1,515 were uniquely annotated as *GT1*s, and 4,037 genes were shared between the two categories.

Since the general HMM models in PFAM and CAZyme are based on protein domains from various species, we aimed to develop a model specifically tailored for *V. vinifera* to identify with precision *GT1*s in *V. vinifera*. GT1s were clustered per protein size and manually curated prior to HMM model generation. Representative GT1s were selected per cluster using sequence similarity and phylogeny. As a result, we identified a total of 4,205 GT1s across the eight *V. vinifera* accessions.

To annotate *GT1* genes in Pixie, we searched the newly assembled Pixie reference for homologs of the 4,205 grapevine *GT1* genes. This yielded 514 *GT1* genes, with 263 in hap1 and 241 in hap2. Their distribution is relatively even across chromosomes (**Figure S7**), with Chromosome 5 containing the highest number (37 in haplotype 1, 36 in haplotype 2), followed by chromosome 18 (26 in haplotype 1, 33 in haplotype 2). Most Pixie *GT1*s (∼95%) were classified into families based on similarity to the 133 characterized *A. thaliana UGTs* (**Figure S8**). A subset that does not match any known family may belong to uncharacterized families or lie outside the *UGT* superfamily.

### A comprehensive Pixie *GT1* expression atlas reveals tissue- and stage-specific patterns of expression

To investigate *GT1* gene expression in Pixie, we built an expression atlas covering roots, stems, buds, flowers, leaves (young, green, senescent), and berries (green, post-veraison, harvest). Three replicates per tissue (30 samples in total) each produced ∼39.51 ± 7.82 million reads. Euclidean distance analysis (**Figure S9A**) confirmed strong replicate reproducibility and identified two main clusters: one encompassing roots, buds, young leaves, flowers, and stems, and another consisting primarily of berry samples and senescent leaves. Principal Component Analysis (PCA) corroborated these clusters, with harvest/post-harvest berry samples forming an additional subset that explained 36% of the variance along PC1 (**Figure S9B**).

Hierarchical clustering (complete linkage) revealed six clusters among the 514 annotated *GT1* genes (**Figure S9C**). Cluster C1 included 32 *GT1* genes that were consistently highly expressed across all tested tissues, suggesting a set of constitutively active genes. Other clusters showed more specialized patterns: C2 genes (10 *GT1*s) were predominantly expressed in roots, while C4 genes (32 *GT1*s) were elevated in roots as well as flowers and buds. Cluster C6 encompassed 7 genes with strong expression in berry tissues during post-veraison stages (harvest and post-harvest), and C7 55 genes upregulated in green and senescent leaves. These tissue-specific expression patterns underscore the diverse functional roles that *GT1* genes may play.

### Baseline *GT1* expression in Pinot Noir berries reveals predominantly uniform patterns with subset-specific variations across ripening stages

Before examining *GT1* expression under smoke exposure, we first investigated their baseline expression across ripening stages. We used a published Pinot Noir RNA-seq dataset (Fasoli et al., 2019) spanning three years, 11 time points, and three berry development stages (pre-veraison, harvest and post-harvest). The transcriptome was strongly influenced by the ripening process, with 57% of the variance explained by time points (**Figure 1A**) making this dataset ideal for our analysis. Additionally, this dataset provides high sequencing depth, frequent sampling intervals, and a close genetic match to Pixie. For this analysis, we constructed a diploid Pinot Noir reference (**Table S1**) and annotated 565 *GT1* genes (240 on hap1, 225 on hap2) using the same workflow as Pixie. The *GT1* distribution closely mirrored Pixie, reflecting their clonal relationship and genomic collinearity (**Figure S10**). After filtering out low-expressed *GT1* genes (TPM < 1), hierarchical clustering on the remaining dataset revealed two main clusters aligned with pre-veraison (tp00– tp04) and post-veraison (tp05–tp11) stages (**Figure 1B, top**).

**Fig. 1.**
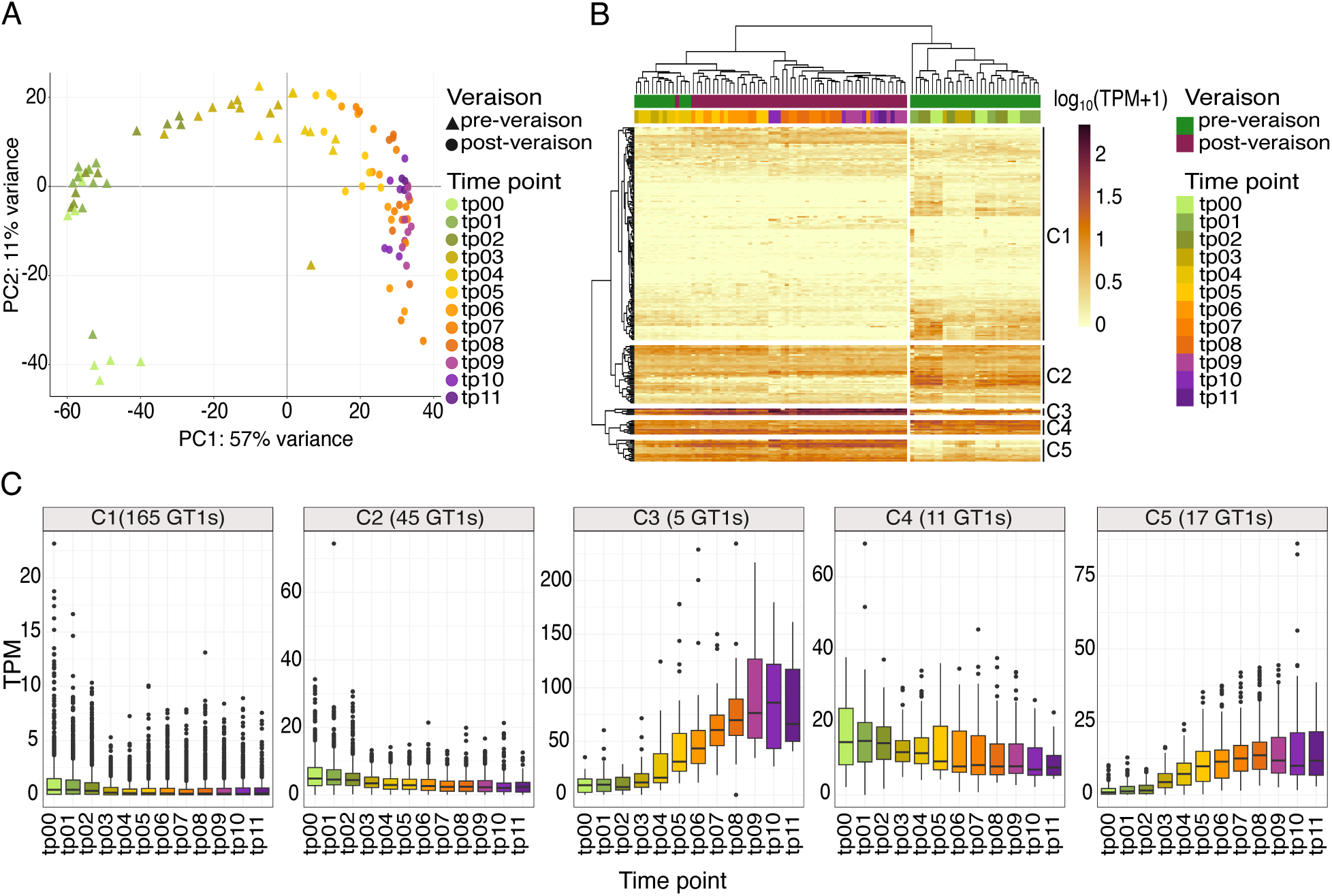
*GT1* expression in grapes during the ripening process. (A) Principal Component Analysis (PCA) of berry transcriptomes expression levels per sample throughout the ripening process. Samples are colored on a gradient from green to purple, representing time points from tp00 (Time Point 0) to tp11 (Time Point 11). Pre-veraison samples are shown as triangles, and post-veraison samples are shown as circles. (B) Heatmap of annotated *GT1* expression (log_10_(TPM+1)) in the Pinot Noir genome. Samples are displayed in columns, with color-coded annotations above indicating sample characteristics: pre-veraison in green, post-veraison in purple. The time point colors align with the PCA in (A). Rows represent individual *GT1s*, clustered hierarchically into five clusters, labeled on the right side of the heatmap. (C) Expression profiles of the Five *GT1* clusters identified in (B) during the ripening process, illustrating distinct patterns of *GT1* expression (in TPM) across time points and developmental stages. Boxplots’s horizontal lines correspond to the 25th, 50th and 75th percentiles, vertical lines extend between the smallest and largest value no further than 1.5 x interquartile range (IQR). Circles beyond the vertical lines represent outliers and extreme values.

GT expression patterns during ripening (**Figure 1B**) were grouped into five clusters (C1–C5) (**Figures 1B and 1C**). Most *GTs* (C1 and C2) showed low, stable expression: C1 (165 GTs) at 0.65 ± 0.001 TPM, and C2 (45 GTs) at 4.03 ± 0.006 TPM but declining after tp02. The smallest cluster, C3 (5 GTs), had the highest post-veraison increase (45 ± 1.96 TPM). Cluster C4 (11 GTs) gradually decreased yet remained above C1 and C2 (12.8 ± 0.24 TPM). C5 (17 GTs) exhibited a pronounced post-veraison rise (9.26 ± 0.23 TPM).

### Smoke exposure induces transcriptional reprogramming and activates glycosylation in ripening grapevine berries

A critical stage for smoke taint formation occurs about seven days post-veraison (Kennison *et al*., 2008). To investigate *GT1* regulation and activity during this period, we exposed Pixie grape clusters to smoke in a bench chamber (**Figure 2A**), conducting two small-scale trials (experiments A and B). Control and smoke-exposed samples were collected one day (“T1”) and three days (“T3”) post-exposure. In experiment B, plants were grown in different greenhouses, introducing additional biological replicates into the results. Experiment B largely corroborated experiment A’s findings. The main text focuses on experiment A, while experiment B results and reproducibility assessments are provided in **Figure S11**.

**Fig. 2.**
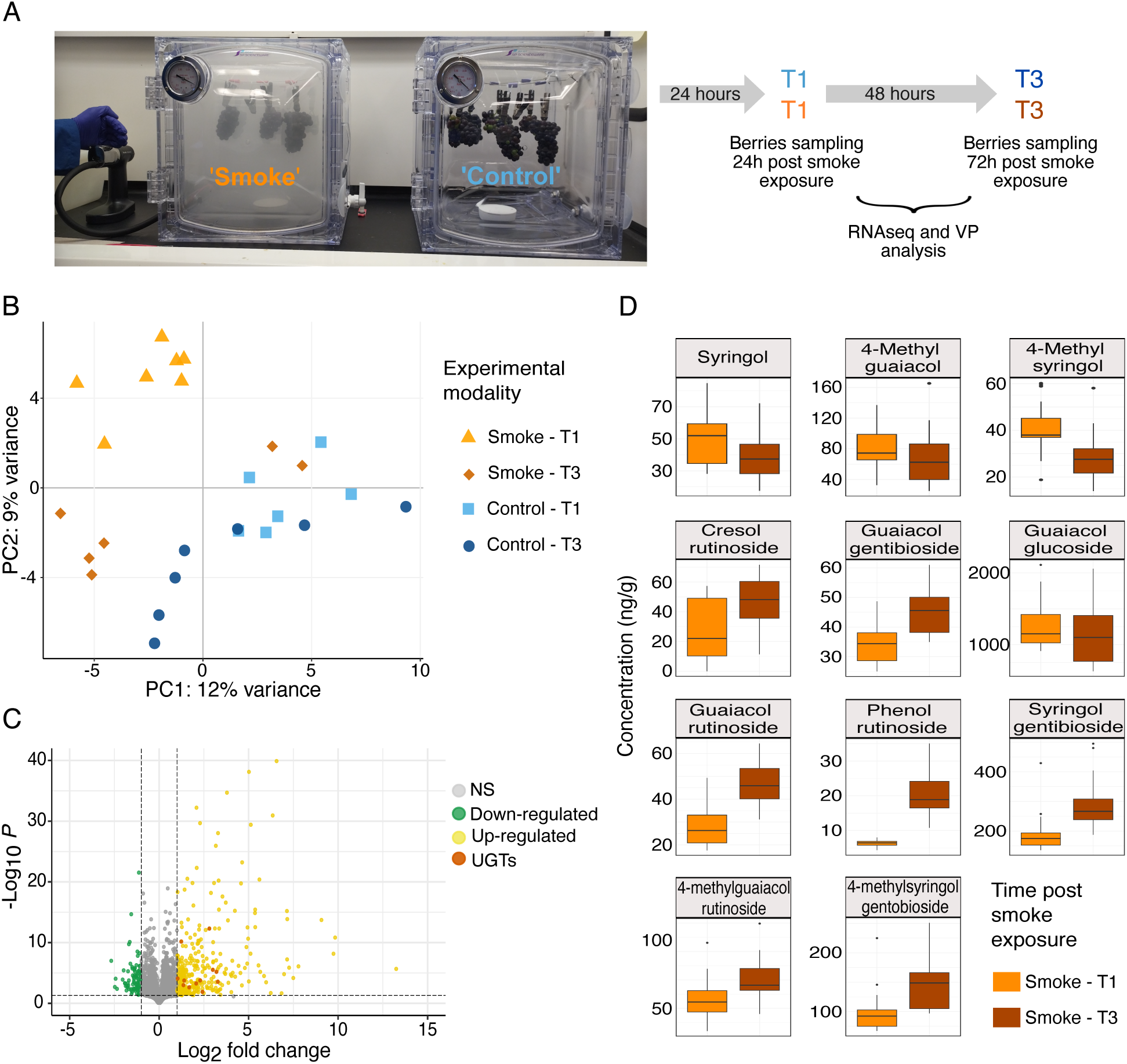
Controlled smoke exposure experiments. (A) Experimental design for the smoke exposure experiment. Two bench-top smoke chambers were used, with Pixie grape clusters exposed to high-concentration smoke in the left chamber (labeled as ’smoke’) and those in the right chamber remaining unexposed (labeled as ’control’). RNAseq, volatile phenols (VPs), and their glycosylated conjugates (Gly-VPs) were analyzed from berries sampled at two time points after exposure: 24 hours post-exposure (’T1’) and 72 hours post-exposure (’T3’). (B) Principal Component Analysis (PCA) of transcriptome expression levels for each sample. Control samples are color-coded in blue, while ’smoke’ samples are color-coded in orange. Samples collected at T1 are represented by circles, and those at T3 are represented by triangles. (C) Volcano plot showing differentially expressed genes (DEGs) in response to smoke exposure. Each point represents a gene, with the x-axis showing log2 fold change and the y-axis showing -log10(P-value). Genes with an adjusted P-value < 0.05 and a log2 fold change > 1 are classified as upregulated (yellow), and those with log2 fold change < -1 as downregulated (blue). Differentially expressed *UGTs* are highlighted in orange. (D) Boxplots of VPs (first row) and Gly-VPs (row n. 2, 3 and 4) showing concentrations (in ng/g and µg/L, respectively) for the smoke-exposed samples at T1 (orange) and T3 (brown). Only VPs with significant differences in concentration (P < 0.05) based on a one-way ANOVA test are displayed. Boxplots’s horizontal lines correspond to the 25th, 50th and 75th percentiles, vertical lines extend between the smallest and largest value no further than 1.5 x interquartile range (IQR). Circles beyond the vertical lines represent outliers and extreme values.

RNA-seq was used to profile the berry transcriptome. Before examining *GT1* expression, we assessed the impact of both smoke and time after exposure on overall transcript profiles. A PCA of the entire transcriptome separated “smoke” samples from “control” samples, with time after exposure also influencing clustering (**Figure 2B**). Comparing “smoke” vs. “control” identified 1,243 DEGs (792 up- and 451 downregulated; **Figure 2C**). GO analysis of upregulated genes highlighted metabolic and stress-response pathways (e.g., catabolic and carbohydrate metabolism, secondary metabolism), while downregulated genes were mostly linked to carbohydrate and primary metabolism, plus more general metabolic and stress responses (**Figure S12A, 12B)** The timing after smoke exposure significantly influenced gene expression, prompting us to analyze the differentially expressed genes (DEGs) between control and smoke-treated samples at T1 and T3 separately. At T1, 379 genes were upregulated, compared to 296 at T3, with 107 genes commonly upregulated at both time points. Conversely, only 123 genes were downregulated at T1 and 138 at T3, with just 7 genes being commonly downregulated at both time points. Upregulated genes at T1 were significantly enriched in biological activities related to stress responses, stimulus responses, catabolic and metabolic processes, and cell death (**Figure S12C, 12D**). At T3, upregulated genes were enriched in regulatory and catalytic activities, secondary metabolic processes, and stress responses (**Figure S12E, 12F**). Overall, these results suggest a significant metabolic and stress-related reprogramming triggered by smoke exposure.

To confirm and assess the impact of smoke on berry chemistry, smoke-derived VP and their glycosylated conjugates (Gly-VPs) were measured. Discernible amounts of VPs (i.e. > 200 ng/g guaiacol) and Gly-VPs were detected in the “smoke” samples compared to those of the “control” ones, as these compounds are often used as biomarkers for smoke exposure (Parker *et al*., 2023). While the VP and Gly-VP levels observed in this study surpassed those found in previously smoke-exposed grape samples likely reflecting the experimental setting (Crews *et al*., 2022*a*; Jiang *et al*., 2022; Parker *et al*., 2023), the primary objective was to identify the transcripts involved in the glycosylation of VPs in grapes. For all the control samples, the concentrations of these biomarker compounds were either below or close to the limit of quantification, so the focus of the results falls on the variations of smoke samples between ‘T1’ and ‘T3’. Three out of 11 measured VPs were found to decrease in concentration from ‘T1’ to ‘T3’ (P value < 0.05), namely syringol, 4- methylguaiacol, and 4-methylsyringol (**Figure 2D**). The concentrations of all 8 measured Gly-VPs increased from T1 to T3 (P value < 0.05). The increment indicated *GT1* activities of transforming the VPs into their glycosylated conjugates. There were no direct correlations observed between the amounts of VPs and their respective glycosides, i.e., guaiacol and guaiacol gentiobioside, as glycosides analyzed in this study were only a small part of all the possible glycosylated products.

### Smoke-inducible UGT genes as potential contributors to smoke taint accumulation in grapevine berries

Among the 512 *GT1*s in Pixie, 12 *GT1* genes were significantly upregulated under smoke exposure (log2 fold changes 1.14–3.41). Notably, smoke exposure did not lead to the downregulation of any *GT1* genes (**Figure 2C**; **Table2**). All 12 smoke-induced *GT1*s belonged to the UGT family and were therefore named *V. vinifera Smoke Inducible UGT1-12* (*VviSIUGT1-12*). They span six UGT families: one UGT71, two UGT72s, one UGT73, six UGT74s (including three pairs from both haplotypes), one UGT90, and one UGT91 (**Table 2**). Structural comparisons with representative *A. thaliana UGTs* with known crystal structure confirmed the highly conserved PSPG box, suggesting these smoke-induced *UGTs* play roles in stress-related detoxification (**Figure S13)**. Hierarchical clustering of the expression profiles of the twelve *VviSIUGTs* (**Figure 3A**) defined two groups: one predominantly comprising control samples at both T1 and T3, plus four smoke-exposed T3 samples, indicating that the initial upregulation subsides by three days post-exposure; and a second cluster exclusively featuring smoke-exposed T1 samples, reflecting an immediate transcriptional response to smoke.

**Table 2:** Pixie smoke-inducible UGT genes.

| Pixie smoke-inducible genes | UGT ID | UGT family* | RNAseq fold-change<br>T1 vs. control, Log2 | qRT-PCR fold-change<br>T1 vs. control, Log2 | Ripening berry expression cluster** |
| --- | --- | --- | --- | --- | --- |
| VITVvi_vPixieFPS01_v1.0.Hap1.chr03.ver1.0.g034780.t01 | *VviSIUGT1* | UGT71 | 1.15 | 1.66 | C1 |
| VITVvi_vPixieFPS01_v1.0.Hap1.chr05.ver1.0.g058910.t01 | *VviSIUGT2* | UGT74 | 2.9 | 2.5 | C1 |
| VITVvi_vPixieFPS01_v1.0.Hap1.chr05.ver1.0.g058920.t01 | *VviSIUGT3* | UGT74 | 3.2 | nd | C1 |
| VITVvi_vPixieFPS01_v1.0.Hap1.chr05.ver1.0.g059090.t01 | *VviSIUGT4* | UGT74 | 3.15 | 2.82 | C1 |
| VITVvi_vPixieFPS01_v1.0.Hap1.chr08.ver1.0.g105080.t01 | *VviSIUGT5* | UGT73 | 1.98 | nd | C1 |
| VITVvi_vPixieFPS01_v1.0.Hap1.chr15.ver1.0.g187030.t01 | *VviSIUGT6* | UGT72 | 1.57 | 1.82 | C1 |
| VITVvi_vPixieFPS01_v1.0.Hap1.chr15.ver1.0.g191090.t01 | *VviSIUGT7* | UGT72 | 1.29 | 1.37 | C3 |
| VITVvi_vPixieFPS01_v1.0.Hap2.chr05.ver1.0.g304320.t01 | *VviSIUGT8* | UGT74 | 1.57 | nd | C1 |
| VITVvi_vPixieFPS01_v1.0.Hap2.chr05.ver1.0.g304330.t01 | *VviSIUGT9* | UGT74 | 3.55 | 2.38 | C1 |
| VITVvi_vPixieFPS01_v1.0.Hap2.chr05.ver1.0.g304520.t01 | *VviSIUGT10* | UGT74 | 2.59 | 3.48 | C1 |
| VITVvi_vPixieFPS01_v1.0.Hap2.chr15.ver1.0.g437150.t01 | *VviSIUGT11* | UGT91 | 2.4 | 1.56 | C3 |
| VITVvi_vPixieFPS01_v1.0.Hap2.chr17.ver1.0.g457760.t01 | *VviSIUGT12* | UGT90 | 3.41 | nd | C1 |

nd: not determined
\* UGT72B1 and UGT74F2 were the *Arabidopsis thaliana* proteins used for structural analysis of the *VviSIUGTs* belonging to UGT72, and UGT74 families, respectively (**Figure S11**).
\*\* *GTL* expression clusters as shown in **Figure 1c**.

**Fig, 3.**
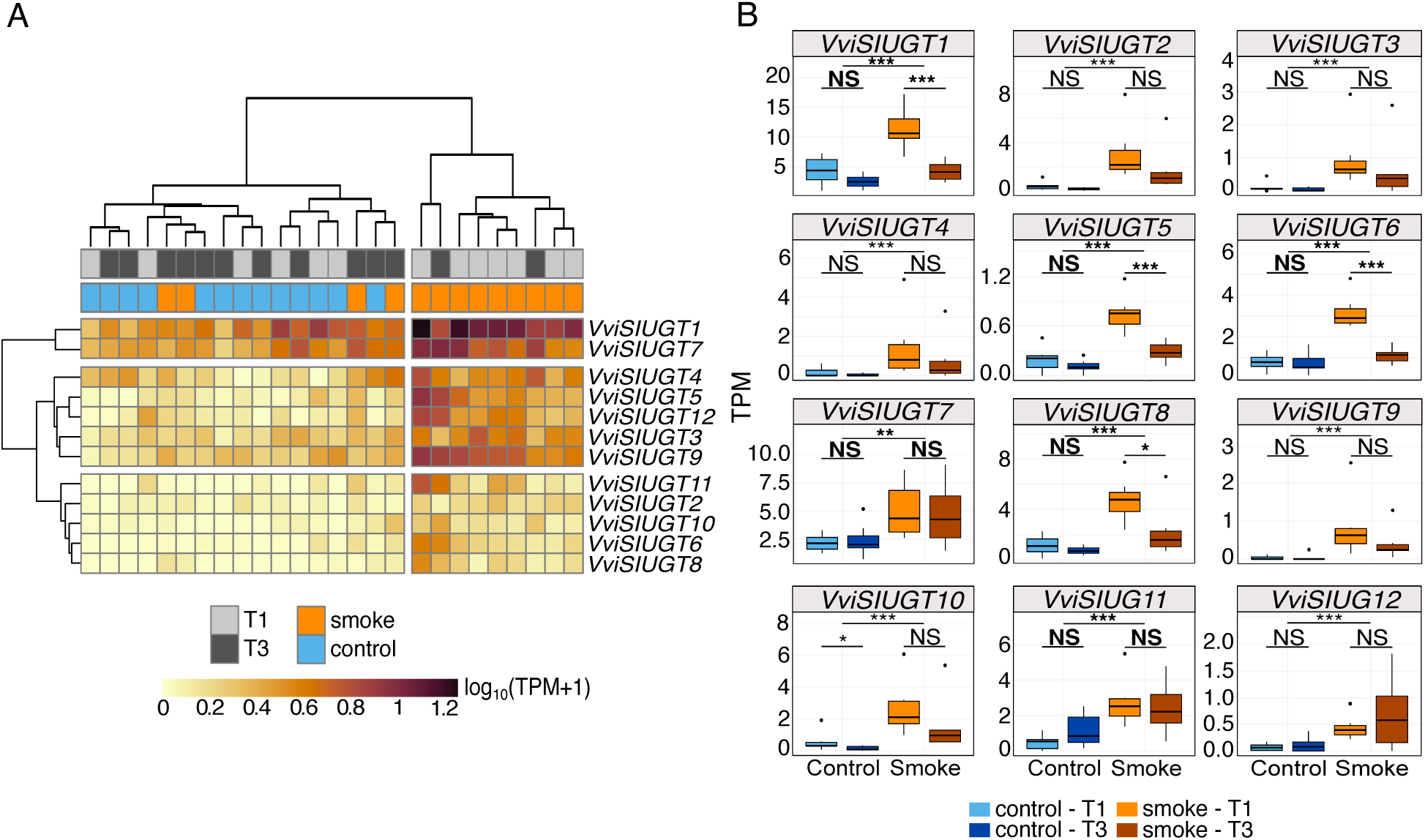
Expression levels of the 12 upregulated *UGTs*. (A) Heatmap showing the expression patterns (log_10_(TPM+1)) of the 12 *UGTs* (rows) across 26 samples (columns). T1 samples are indicated in light gray, T3 samples in dark gray, control samples in blue, and smoke samples in orange at the top of the heatmap. Both *UGTs* (rows) and samples (columns) were clustered using hierarchical clustering with the complete linkage method (hclust), represented by dendrograms at the top (samples) and left (*UGTs*) of the matrix. (B) Boxplots display the expression levels of each UGT, with control/T1 in light blue, control/T3 in dark blue, smoke/T1 in orange, and smoke/T3 in dark red. Significant differences are indicated above each boxplot: (NS) for non-significant, (**) for P < 0.01, and (***) for P < 0.001. A two-way ANOVA was used for genes with normally distributed data (highlighted in bold: VviSIU*GT1*-5-6-7-8-11), while a Wilcoxon test was used for non-normally distributed data. Boxplots’s horizontal lines correspond to the 25th, 50th and 75th percentiles, vertical lines extend between the smallest and largest value no further than 1.5 x interquartile range (IQR). Circles beyond the vertical lines represent outliers and extreme values.

In smoke-treated samples, most of the 12 *UGTs* trended toward a return to basal expression levels at T3 with four exhibiting a significant decline compared to the T1 timepoint (Wilcoxon test, *P*- value < 0.01) (**Figure 3B**); notably, these *UGTs* showed no significant differences from the control at T3, indicating a transient spike in UGT expression 24 hours following smoke exposure. A second, independent trial (Experiment B, **Figure S9**) confirmed these patterns: 9 *UGTs* were significantly upregulated, with none downregulated. Four of these *UGTs* (*VviSIUGT1*, *VviSIUGT6*, *VviSIUGT10*, *VviSIUGT11*) were consistently upregulated by smoke exposure in both experiments with elevated expression at T1 relative to T3. qRT-PCR analysis was performed for eight selected *VviSIUGT* genes using three biologically replicated RNA samples. The results show a significant induction for each transcript 24 hours (T1) after smoke exposure (**Table 2, Figure S14**). The induced fold changes calculated from the RNA-seq and qRT-PCR datasets were consistent with correlation coefficients ranging from 0.68 to 1.0 (Figure S14).

Beyond smoke conditions, these *VviSIUGTs* also showed variability across tissues in the Pixie expression atlas, with the highest expression typically seen in senescent leaves, followed by flowers and green leaves (**Figure S15**). Among them, *VviSIUGT7* stood out for consistently elevated expression in roots (4,276.20 TPM), senescent leaves (3,841.16 TPM), and adult leaves (2,740.94 TPM), and it showed higher expression levels before veraison in Pixie berries. In contrast, the corresponding 12 *VviSIUGTs* in Pinot Noir remained largely stable during berry ripening: most grouped into a minimally variable cluster (C1), while *VviSIUGT7* and *VviSIUGT11* displayed moderate variation between pre- and post-veraison stages (C3) (**Figure S16**).

Collectively, these findings demonstrated that the 12 identified *VviSIUGTs* respond robustly yet transiently to smoke exposure while also exhibiting notable tissue- and developmental-stage specificity. Given their plausible roles in smoke taint formation, we focused our subsequent analyses on elucidating their regulation.

### Co-expression network analysis reveals gene modules linked to candidate *UGTs* and smoke exposure

We performed a co-expression network analysis to identify genes that may respond similarly to smoke exposure as the twelve *VviSIUGTs*. Using the WGCNA R package (Langfelder and Horvath, 2008), we detected 83 gene modules based on co-expression patterns (**Figure S17**). We then examined the correlations of these modules with our experimental factors (“smoke” vs. “control,” “T1” vs. “T3”) and with free and bound volatile phenols (**Figure S18**). Eight modules exhibited a positive correlation above 0.4 with smoke exposure (p < 0.05): Paleturquoise (corr = 0.70), Midnightblue (corr = 0.57), Lightgreen (corr = 0.49), Thistle3 (corr = 0.48), Purple (corr = 0.45), Pink and Antiquewhite (corr = 0.43), and Tan (corr = 0.40). We evaluated the Module Membership (MM)—which measures the correlation of a gene to its module eigengene—to pinpoint where each VviUGT mapped, revealing that all 12 *VviSIUGTs* clustered in four smoke- correlated modules: Purple, Midnightblue, Lightgreen, and Paleturquoise (**Figure 4A, 4B**).

**Fig. 4.**
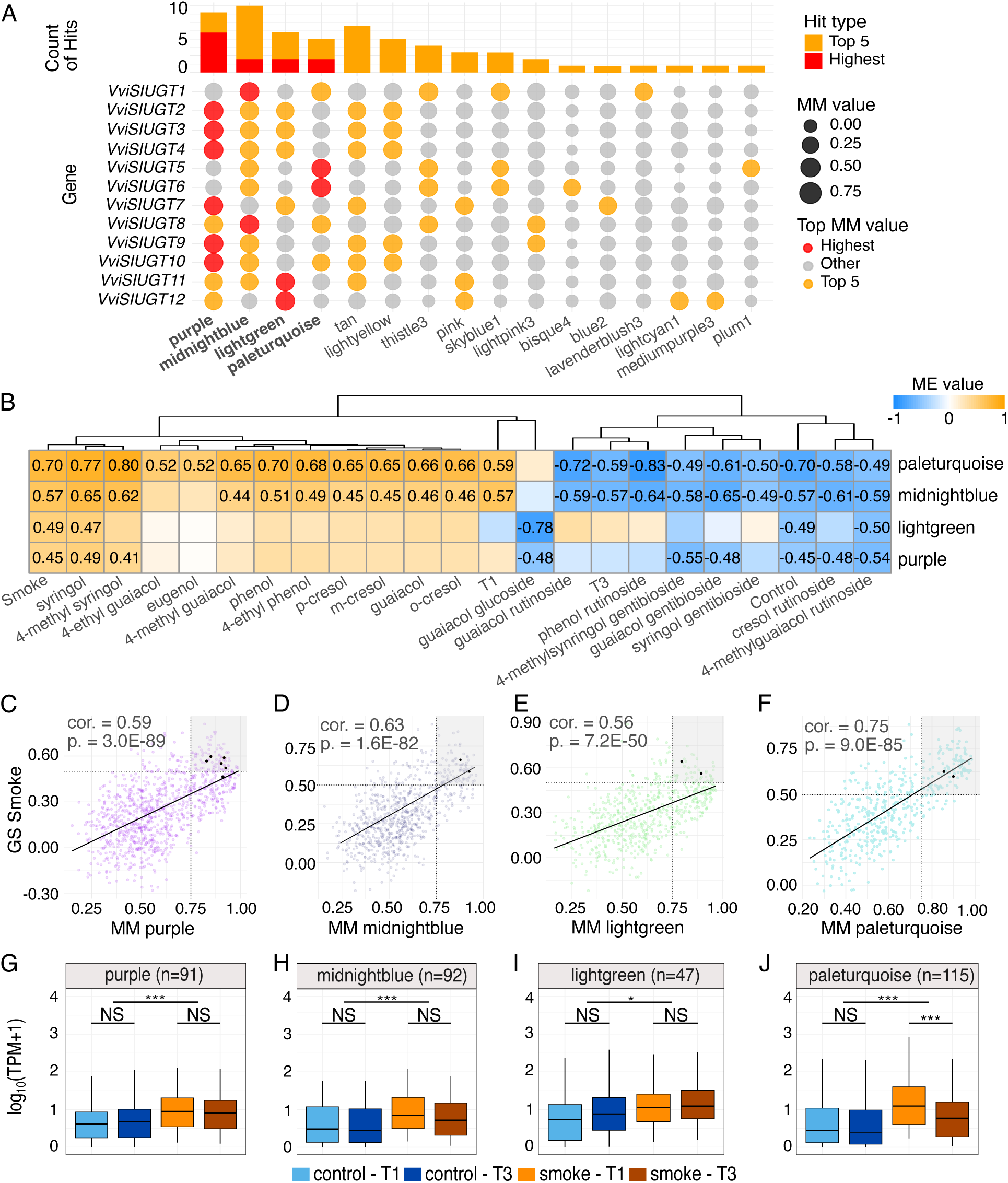
Gene Co-expression Network Analysis. (A) Module Membership (MM) of 12 candidate *UGTs* across WGCNA modules (columns). Each UGT (row) is represented by a gray circle, with the circle size reflecting the MM value: the larger the circle, the higher the MM value for that gene within the corresponding module. The highest MM value per gene (referred to as a “hit”) is highlighted in orange, with the overall highest MM value emphasized in red. The top bar plot shows the count of hits per module, prioritizing the highest hit per gene, followed by the top 5. The four best-performing modules are highlighted in bold. (B) Heatmap of Correlation Between Selected Module Eigen-genes and Experimental Traits (Modality, Time (T), and Volatile Phenols (VPs)). Each cell color represents the Module Eigen-genes value, the Pearson correlation coefficient is displayed in each cell with significant correlations (P < 0.05). The top dendrogram indicates the hierarchical clustering of the traits based on all modules (a complete module-trait correlation heatmap is available in the supplementary materials). (C, D, E, F) Correlation between MM and Gene Significance (GS) for the smoke trait for each of the four selected modules—purple (C), midnightblue (D), lightgreen (E), and paleturquoise (F). The black diagonal line represents the regression line with the correlation (cor.) and *p*-value (p.) indicated in the top left corner of each plot. Dashed lines represent thresholds of GS > 0.5 and MM > 0.75, helping to identify hub genes in the top right corner of each plot, highlighted in gray. The 12 candidate GTs are highlighted in black. (G, H, I, J) Boxplots representing the expression levels (log10(TPM+1) for comparison purposes) of the hub genes in each of the four selected modules—purple (G), midnightblue (H), lightgreen (I), and paleturquoise (J)—across experimental conditions: control/T1 (light blue), control/T3 (dark blue), smoke/T1 (orange), and smoke/T3 (dark red). Significant differences based on a two-way ANOVA are indicated above each boxplot: (NS) for non-significant, () for P < 0.05, and (**) for P < 0.001. The total number of hub genes for each module is noted as “n”. Boxplots’s horizontal lines correspond to the 25th, 50th and 75th percentiles, vertical lines extend between the smallest and largest value no further than 1.5 x interquartile range (IQR).

Notably, all measured VPs correlate positively with the four modules, while most Gly-VPs correlate negatively (**Figure 4B**). This pattern likely reflects glycosylation’s role in detoxifying grape berries from xenobiotics like smoke-related VPs. Excessive VPs trigger smoke-correlated modules, while Gly-VP accumulation signals response termination (Hewitt *et al*., 2024). A positive relationship (P < 0.001) between MM and Gene Significance (GS) indicated that genes central to these four modules are strongly associated with smoke exposure (**Figure 4C, 4F**).

We then identified hub gene (i.e., highly connected genes within a network that may play central regulatory roles (MM > 0.75 and GS > 0.5)), within each module uncovering 91 hubs in Purple, 92 in Midnightblue, 47 in Lightgreen, and 116 in Paleturquoise. Interestingly, all but one of the *VviSIUGTs* appeared among these hubs (**Figure 4C, 4F**). To characterize the genes co-expressed with the *VviSIUGTs*, we examined each module’s hub genes in detail (**Figure S19**, **Table S4**). In Purple, eight chalcone synthase genes (all on chromosome 16) and two high-MM glutathione S-transferases (GSTs) were linked to secondary metabolism and stress responses; additional TFs (e.g., ABR1-like, WRKYs) pointed to stress-adaptive roles. Midnightblue lacked significant GO enrichments but included highly smoke-correlated genes for sugar metabolism (sucrase/ferredoxin family, GS=0.87) and flavonoid biosynthesis (naringenin 2-oxoglutarate 3-dioxygenase, GS=0.84). Its top-MM genes, such as a drug-resistance ABC transporter, similarly suggested stress-related secondary metabolism. Lightgreen showed enrichment (Padj < 0.05) in stress and defense processes, with defense regulators (NDR1/HIN1-like, WRKY) and a carboxylesterase (GS=0.75), suggesting a detoxification link to smoke exposure. Paleturquoise was enriched (Padj < 0.005) in secondary metabolic processes, responses to chemicals, and stimuli; it contained multiple GSTs, mannitol dehydrogenases (MDHs), and four naringenin 2-oxoglutarate 3- dioxygenases (F3Hs). High-GS F3Hs and top-MM GSTs reinforce a role in stress tolerance and flavonoid pathways.

When we compared hub-gene expression levels (log10(TPM+1)) between ‘smoke’ and ‘control’ as well as T1 vs. T3 (**Fig. 4G–J**), we observed significantly higher expression under smoke in all four modules, mirroring the behavior of *VviSIUGTs*. Only the Paleturquoise module showed a drop from T1 to T3 in smoke, matching the transient spike seen in several of the VviSI*UGTs*. Collectively, these results highlight that the *VviSIUGTs* reside in modules (Purple, Midnightblue, Lightgreen, Paleturquoise) strongly correlated with smoke exposure and enriched in genes involved in secondary metabolism, stress defense, and transcriptional regulation. To further elucidate how these co-expressed networks are regulated under smoke stress, we investigated potential enriched transcription factor binding sites in the promoters of hub genes, seeking regulatory signals that could be responsible for the rapid but transient upregulation of the *VviSIUGTs*.

### Transcription factor binding site analysis identified stress-responsive elements in smoke-responsive modules

To explore the regulatory mechanisms underlying expression of the smoke-responsive *VviUGTs* and their co-expressed genes, we examined the transcription factor binding sites (TFBS) present in the 3-kb upstream regions (relative to the transcription start site) of the hub genes in each of the four modules. We found several TFBS significantly enriched, indicating potential regulators of these co-expressed genes under smoke conditions.

Among the 12 candidate *VviSIUGTs*, five candidates carried one or more enriched TFBS in their promoter region. The candidates *VviUGT3, VviUGT7, VviUGT9,* and *VviUGT10*, associated with the purple module, carried the WRKY8 TFBS. In the purple module, the WRKY8 TFBS (adjusted P-value = 0.0128) was enriched in 45 of the 91 hub genes, including the four of the six candidate *UGTs* associated with this module: 33 of these were upregulated under smoke, while 12 showed no changes. Among the purple hub genes, eight were annotated as WRKY transcription factors, including two hub TFs that were not smoke-induced but were actively expressed. Across the genome, 166 WRKYs were identified, with 27 showing module membership (MM) > 0.50 in the purple module.

*VviUGT5,* associated with the paleturquoise module, carried four enriched sites: NAC002, BZIP60, TGA10 and TGA3. In the paleturquoise module, nine TFBS were enriched (adjusted P-value < 0.05), including NAC002 from the CGM class, four bZIP-type sites (bZIP60, bZIP910, PK09021.1, ZmbZIP54), and four TGA-type sites (TGA10, TGA2, TGA3, TGA7). Among the 116 hub genes, 101 contained at least one of these TFBS and 57 were upregulated by smoke. Although nine hub genes were annotated as TFs, none were differentially expressed. Ten NAC and four bZIP TFs, each with MM > 0.50, were also identified as potential regulators within paleturquoise.

In the two other modules, no significantly enriched TFSB were present in the candidate *VviSIUGTs*, however, we found interesting enriched TFBS in their co-expressed genes. In the midnightblue module, the TFBS Zm00001d015407 (adjusted P-value = 0.015) was found in 11 hub genes, eight of which were smoke-upregulated. One hub gene was annotated as a TF (HBP- 1b(c1)) but was not differentially expressed. Midnightblue contained 19 TF-annotated transcripts, three of which were upregulated by smoke. A probable WRKY75 (*VITVvi_vPixieFPS01_v1.0.Hap1.chr14.ver1.0.g178980.t01*) with MM = 0.57 showed partial overlap with the purple module. No significant TFBS enrichment was detected in the lightgreen module. Nonetheless, 23 hub transcripts were annotated as TFs, with three upregulated under smoke (two AP2-domain factors and one probable WRKY factor, *VITVvi_vPixieFPS01_v1.0.Hap1.chr19.ver1.0.g239180.t01*, MM = 0.78 in the purple module). Three additional WRKY TFs were identified in lightgreen, though they were not differentially expressed under smoke conditions.

Collectively, these results indicate that multiple TFs, particularly WRKY, NAC, bZIP, and TGA, are potential regulators of the smoke-responsive *VviUGTs* and their co-expressed genes. While few TFs in these modules were themselves differentially expressed under smoke, their prominent module memberships and enriched TFBS in hub-gene promoters point to a coordinated regulatory network that may drive both the immediate and sustained responses to smoke exposure.

## Discussion

To identify candidate genes involved in smoke taint in wine, we conducted a controlled smoke exposure experiment. We hypothesized that enzymes from the GT1 family play a role in the accumulation of smoke-derived compounds associated with smoke taint. Our analysis focused on *GT1* gene expression and VP dynamics during smoke exposure. Using the dwarf grapevine Pixie as a model system, we identified 12 candidate UGTs potentially linked to smoke taint accumulation in grape. Quantification of both VP and gly-VP levels in berries during exposure revealed a strong correlation between the upregulation of *GT1* genes and VP accumulation. Additionally, we identified genes co-expressed with the 12 candidate UGTs. Promoter analysis of these co-expressed genes revealed transcription factor binding sites (TFBS) shared with the candidate UGTs, suggesting a regulatory network potentially contributing to the accumulation of smoke taint.

Using Pixie as a diploid reference genome provides advantages for *in planta* experiments compared to the haploid reference PN40024. Pixie originates from the L1 layer of Pinot Meunier, a chimeric mutation of Pinot Noir and Pinot Gris (Boss and Thomas, 2002; Cousins, 2012). Its clonal relationship with PN40024, widely used in grape research, facilitates cross-referencing and comparative genomics. Pixie’s dwarfism and continuous flowering (Cousins, 2012; Pellegrino *et al*., 2019) make it ideal for controlled experiments, allowing the use of bench-sized smoke chambers for full grape clusters. Year-round flowering eliminates seasonal constraints, enabling experiments from flowering to harvest. Additionally, the Pixie-specific gene expression atlas developed in this study enhances its utility by providing baseline gene expression profiles across tissues and developmental stages.

UDP-glycosyltransferases (UGTs) of the GT1 family play essential roles in plant growth, development, and stress adaptation (Gharabli *et al*., 2023). In *Arabidopsis thaliana*, *UGT79B2* and *UGT79B3* enhance anthocyanin production and abiotic stress tolerance 2/11*/25 8*:22:00 PM, while in rice, overexpression of *GSA1* (*UGT83A1*) improves drought, salt, and heat tolerance (Dong *et al*., 2020). In this study, 12 *V. vinifera* UGT-encoding genes, designated as smoke-induced *UGTs* (*VviSIUGT1–12*), were upregulated in response to smoke exposure, with four (*VviSIUGT1, 6, 10, 11*) consistently upregulated across experiments. Additionally, four candidates (*VviSIUGT1, VviSIUGT2, VviSIUGT4, and VviSIUGT7*) were previously linked to smoke exposure in grapevines (Hewitt *et al*., 2024). Moreover, *UGT72B27*, identified by Härtl et al. (2017) for its role in phenolic glucoside production during smoke exposure, shares close homology with *VviSIUGT7*, further supporting its functional relevance.

Co-expression analysis revealed key stress response and smoke adaptation mechanisms. Our 12 candidate genes clustered into four networks enriched in detoxification, stress defense, secondary metabolism, and transcriptional regulation, highlighting their role in environmental adaptability. Moreover, all but one of them are considered hub genes in their respective networks, emphasizing their role in response to smoke exposure. Glycosylation is crucial for detoxifying xenobiotics, including smoke-derived phenolics in grapes. Plants employ a sophisticated detoxification system to convert xenobiotics into less toxic forms (Sandermann, 1992; Kawahigashi, 2009). In grapes exposed to wildfire smoke, this mechanism is central to mitigating the effects of smoke-derived phenolic compounds, as these compounds are metabolized into non-volatile glycoconjugates. While this detoxification pathway protects the plant, it poses challenges for human consumption, as these glycoconjugates release volatile phenols during fermentation, wine aging (Kennison *et al*., 2008), and wine consumption (Mayr *et al*., 2014), leading to undesirable smoke taint (Hayasaka *et al*., 2010; Mayr *et al*., 2014; Bilogrevic *et al*., 2023). In our study, six *UGTs* (*VviSIUGT2, 3, 4, 7, 9, 10*) co-expressed with cytochrome P450s and glutathione-S-transferases (GSTs) in the purple module, suggesting a coordinated detoxification role (Cummins *et al*., 2011; Pandian *et al*., 2020), aligning with Hewitt et al. (2024). The paleturquoise module, showing decreased hub gene expression between T1 and T3 and including *VviSIUGT5* and *6*, was enriched in mannitol dehydrogenase and GSTs. ROS signaling is an early stress response in plants (Mittler *et al*., 2022; Ang *et al*., 2024), and these genes are likely to contribute to rapid smoke exposure responses.

Beyond identifying co-expressed genes, we also analyzed the presence of enriched transcription factor binding sites within the hub genes of the co-expression module. Notably, we detected significant enrichment of WRKY, NAC, bZIP, and TGA families, which are regulators of abiotic stress responses (Khoso *et al*., 2022; Han *et al*., 2023; Fuertes-Aguilar and Matilla, 2024; Lu *et al*., 2024). These TFBS were present in the promoter region of five of our candidates (*VviSIUGT3*, *5*, *7*, *9* and *10*). WRKY TFs were particularly prominent in our results. The WRKY TF family, one of the largest in higher plants (Jiang *et al*., 2017), is central to transcriptional reprogramming during plant stress responses (Khoso *et al*., 2022). During smoke exposure, the upregulation of mannitol dehydrogenase, a key player in redox homeostasis and osmotic stress regulation, has been previously reported (Hewitt *et al*., 2024) and is also observed in our results. The involvement of WRKY TFs in regulating such pathways suggests their potential role in mitigating oxidative stress and maintaining cellular homeostasis during smoke exposure (Khoso *et al*., 2022). NAC TFs play critical roles in abiotic stress tolerance (Fuertes-Aguilar and Matilla, 2024), such as cold (Diao *et al*., 2020), drought (Li *et al*., 2023) or salt stress tolerance (Zhang *et al*., 2021). TFs from the bZIP family are implicated in linking environmental cues to secondary metabolite biosynthesis, as seen in tobacco and barley under stress conditions (Collin *et al*., 2020; Duan *et al*., 2022). Finally, TGA TFs, a clade of TFs belonging to the bZIP family, emerged as central players in integrating phytohormone signaling pathways, enhancing tolerance to diverse stresses, such as drought (Zhong *et al*., 2015), salinity (Du *et al*., 2014), and heavy metal exposure (Farinati *et al*., 2010).

To further investigate the role of the 12 *VviSIUGTs*, we compared their transcriptional activity with volatile phenol (VP) concentrations in smoke-exposed berries. While no clear correlation was observed between volatile phenol levels and their glycosylated conjugates, the increase in glycosides from T1 to T3 aligned with peak UGT expression at T1. The rapid uptake of VPs from smoke by grape berries, consistent with previous reports (Szeto *et al*., 2020), was evident within one hour post-exposure. Gly-VP concentrations increased significantly between one day and one week post-exposure, remaining elevated until harvest. Among the analyzed compounds, guaiacol glucoside exhibited higher concentrations than other disaccharides, though this observation requires further validation. Interestingly, guaiacol glucoside, the only monoglycoside analyzed, showed a smaller increase from T1 to T3 compared to the diglycosides, suggesting a preference for diglycoside formation, corroborating previous findings (Dungey *et al*., 2011; Crews *et al*., 2022*b*). These results indicate that GT1 transcriptional activity peaks within a short timeframe (<24 hours), with reduced expression at T3 potentially reflecting the upregulation of other GTs, particularly GT2 and GT3. In tulip flowers (*Tulipa* cv. Apeldoorn), GT2 and GT3 primarily contribute to flavonol diglycoside formation (Kleinehollenhorst *et al*., 1982), whereas GT1 is essential for monoglycoside production, acting as a precursor for GT2- and GT3-mediated glycosylation. A similar mechanism may occur in grape berries, as suggested by the shifting dynamics of GT1 expression and Gly-VP accumulation from T1 to T3.

To build on our findings, several future directions can provide a more comprehensive understanding of the role of GT1 in smoke taint. First, functional validation of the identified candidate UGTs through gene knockout experiments will be essential to confirm their involvement in smoke taint. Additionally, testing the complete process on transformed grapes, from grape exposure to wildfire smoke through winemaking and sensory evaluation, will help establish the direct impact of these genes on smoke taint perception in wine. These future perspectives are crucial for breeding programs focused on selecting grape varieties tolerant to smoke exposure, an increasingly pressing issue due to the rise in uncontrolled wildfires. The rapid transcriptional responses we observed at early time points (between T1 and T3) raise intriguing questions about the kinetics of *GT1* regulation. Conducting experiments with additional, more frequent time points would help determine the precise dynamics of gene expression in response to smoke. Another key consideration is the impact of smoke concentration. The impact of smoke concentration on berry composition varies largely depending on grape cultivar and the smoke-marker compound (Härtl *et al*., 2017). In addition, smoke exposure in the vineyard can vary significantly based on factors like exposure duration, proximity to the fire, and wind direction. Testing multiple concentrations in controlled settings as well as in vineyards would provide valuable insights into the practical implications for vineyard management. While our study focused on berries, we observed that GT1s expression varies across tissues, and the impact of smoke on other grapevine tissues, such as leaves, warrants investigation. Understanding tissue-specific responses to smoke exposure could reveal additional mechanisms influencing the development of smoke taint and inform more targeted mitigation strategies. Finally, although it has been shown that smoke-derived compounds accumulate in the pulp and skin of grape berries, it would be fascinating to explore the impact of wildfire smoke at the cellular level. With advancements in single-cell analysis techniques for plants, applying these methods to study smoke exposure in grape berries could answer critical questions. For instance, which cell types are primarily involved in the development of smoke taint? Does the skin type of the berries, which can vary between grape varieties, influence the absorption of smoke-derived compounds? Such insights could deepen our understanding of the cellular mechanisms behind smoke taint and reveal varietal differences that might affect susceptibility to smoke exposure.

## Acknowledgments

We thank the UC Davis Genome Center DNA Technologies Core Facility for their assistance with sequencing and Dr. Andrea Minio for his bioinformatics support. This research was supported by a USDA Agricultural Research Service cooperative agreement (#58-2030-2-033) and CRIS project 2030–21220–003-000-D. The project was also partially supported by the E. & J. Gallo Winery and the Ray Rossi Endowment in Viticulture and Enology. Mention of trade names or commercial products is solely for the purpose of providing specific information and does not imply a recommendation or endorsement by the U.S. Department of Agriculture. USDA is an equal opportunity provider and employer.

## Notes

### Competing Interest Statement

The authors have declared no competing interest.

